# Precocial ancestry of placental mammals

**DOI:** 10.1101/2025.10.30.685703

**Authors:** Arun R. Chavan, Mihaela Pavlicev, Günter P. Wagner

## Abstract

Mammals exhibit two distinct reproductive strategies: altriciality, where neonates are highly underdeveloped, and precociality, where neonates are well-developed. The neonatal developmental state of deep nodes in mammalian phylogeny remains unresolved. Here, we use phylogenetic comparative methods to reconstruct the evolutionary history of neonatal maturity in mammals and demonstrate that the eutherian ancestor likely gave birth to precocial neonates. This finding contradicts the prevailing view that precociality evolved multiple times within the eutherian lineage. We contextualize this result with three lines of evidence. First, recent fossil evidence suggests that precocial life history traits arose early in therian evolution. Second, altricial eutherian neonates are markedly more developed at birth compared to altricial non-eutherians, suggesting a precocial ancestry in eutherian evolution. Third, reproductive traits that enable prolonged pregnancy originated in the stem lineage prior to the eutherian ancestor. Based on these findings, we propose an alternative model for the evolutionary history of precociality in mammals.

## 2 Introduction

Mammalian neonatal maturity spans a spectrum from altricial to precocial (Portmann, 1945). Altricial neonates (e.g., murine rodents), born highly underdeveloped, are often hairless and unable to thermoregulate or move independently. They typically do not open their eyes for days after birth and require extensive parental care for early survival. In contrast, precocial neonates (e.g., ruminants) are far more developed. They are born with fur, can open their eyes, move independently shortly after birth, and require significantly less parental care.

Although neonatal maturity in mammals exists along a continuum, it follows a notably bimodal pattern as illustrated by the distribution of gestation length relative to body mass (Martin & MacLarnon, 1985; **Supp. Figure 1**). Most species fall distinctly into either the altricial or precocial category, with relatively few occupying the intermediate states. This bimodal pattern suggests that altriciality and precociality represent two distinct reproductive strategies.

Understanding the evolutionary history of neonatal maturity is pivotal for uncovering broader patterns in mammalian life history evolution. The divergence between these reproductive strategies results from selection on a suite of developmental, physiological, and life history traits. These include litter size, growth rate, neonatal body mass relative to the adult (Derrickson, 1992; Grunstra et al., 2019), relative allocation of resources between gestation and lactation (Künkele & Trillmich, 1997), and mating system (Zeveloff & Boyce, 1980). Because these traits are interdependent (Danis et al., 2025), the adoption of an altricial or precocial strategy accordingly influences their evolutionary trajectories.

Evolutionary history of neonatal maturity has been a source of debate not only in mammals but also across other amniotes (Starck & Ricklefs, 1998; Werneburg et al., 2016). Due to the lack of definitive fossil correlates of neonatal maturity, investigation of the evolution of this trait remains speculative. With the exception of a few studies (Ferner et al., 2017; O’Leary et al., 2013; Werneburg et al., 2016), the inference of ancestral states has been predominantly narrative rather than analytical. The prevailing view is that neonates were altricial at deeper nodes in mammalian phylogeny, including the mammalian, therian, and eutherian (placental) ancestors, with precociality evolving multiple times within the eutherian lineage. While the altriciality of the therian ancestor has been extensively debated from contrasting perspectives (Lillegraven, 1975; Lillegraven et al., 1987; Renfree, 1993; Smith, 2001; Smith & Keyte, 2020; White et al., 2023), the altriciality of the eutherian ancestor has received little scrutiny.

Here, we reconstruct the evolutionary history of neonatal maturity using phylogenetic comparative methods with a dataset of 420 mammalian species and a modern phylogenetic framework that accounts for phylogenetic uncertainty. Our results suggest that the eutherian ancestor was precocial rather than altricial. We discuss this result in the context of recent fossil and comparative evidence and propose an alternative model for the evolution of mammalian reproductive mode.

## 3 Results

### 3.1 Inferred ancestral state for eutherian mammals is precocial

Neonatal maturity data for 420 mammals (**Supp. Table 1**) were compiled from Case (1978) and PanTHERIA (Jones et al., 2009). Case (1978) provides detailed information for 150 mammals, but with a taxonomic representation skewed toward primates and other boreoeutherian species. To address this bias, additional data from PanTHERIA were incorporated, producing a more balanced dataset representing 21 of 27 mammalian orders and 16 of 19 eutherian orders (**Supp. Table 2**).

Ancestral state reconstruction was performed using stochastic character mapping (Bollback, 2006; Huelsenbeck et al., 2003; Revell, 2025). Phylogenetic uncertainty was accounted for by using a sample of 100 phylogenetic trees from a distribution of possible trees (Upham et al., 2019) and running ancestral state reconstruction on each. Three models of trait evolution — equal rates (ER), symmetric (SYM), and all rates different (ARD) — were fitted to the data. To account for model uncertainty, 1,000 stochastic character histories were generated using parameters estimated from all three models, with the number of histories per model determined by their respective model weights. Posterior probability of character states at each node was calculated by integrating over the 1,000 stochastic character maps. The same analysis was performed on the consensus tree for visualization.

The ARD model provided the best fit to the data across all 100 sampled trees by a wide margin (**Supp. Figure 2**). The mean model weight for ARD across the trees was 0.989, compared to 0.003 for the ER model and 0.009 for the SYM model. Consequently, the majority of character histories were generated using the ARD model parameters.

As expected, the ancestral states for Prototheria (monotremes) and Metatheria (marsupials) are consistently altricial in all sampled trees, with mean posterior probabilities of 0.93 and 0.94, respectively (**Figure 1**). For Theria and Mammalia, the ancestral states are ambiguous but lean toward altriciality. For the therian ancestor, the mean posterior probabilities are 0.6 for altriciality and 0.36 for precociality, while for the mammalian ancestor, they are 0.69 and 0.28, respectively.

**Figure 1:**
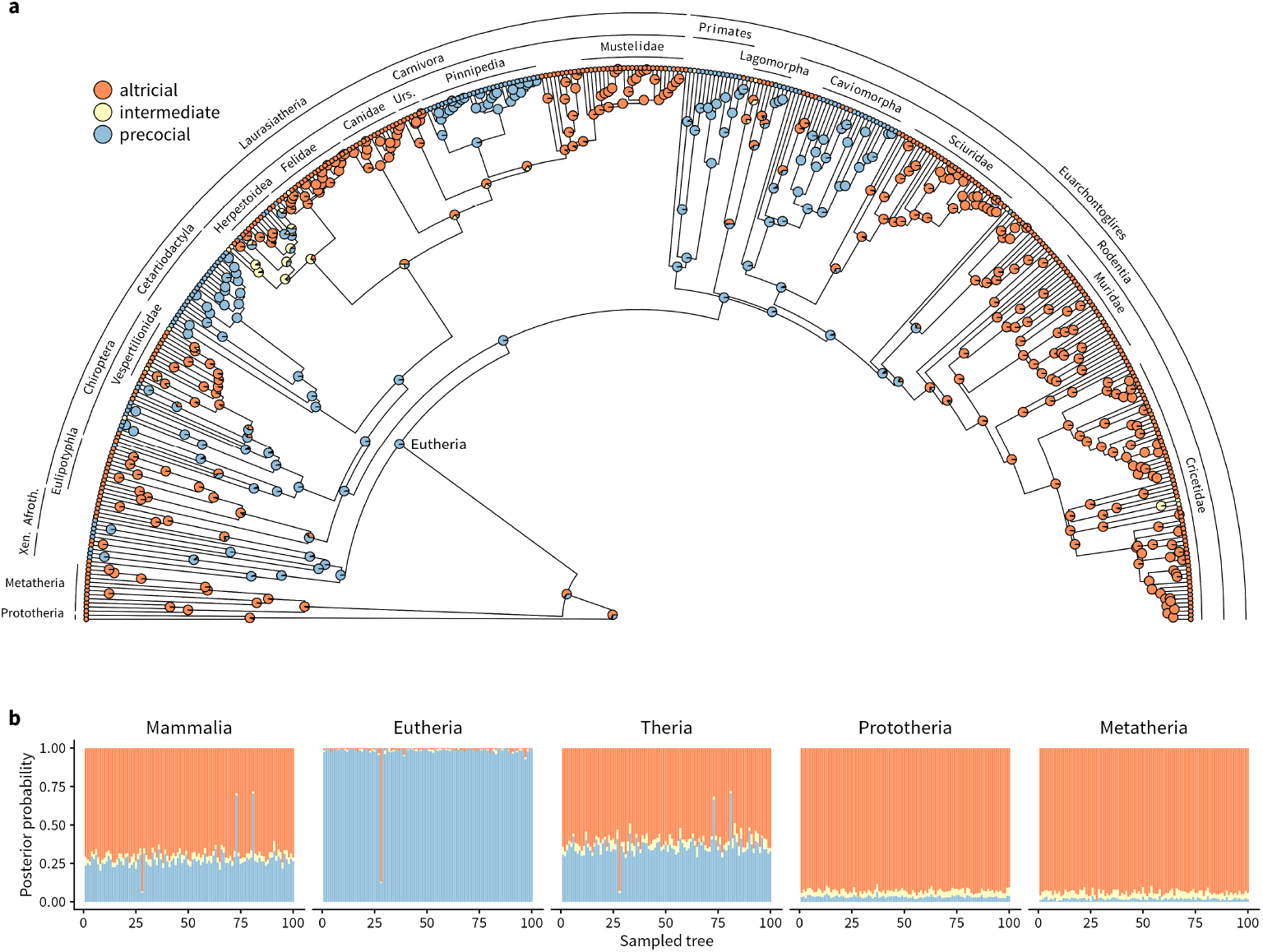
Ancestral state reconstruction of neonatal maturity using stochastic character mapping. **(a)** Consensus phylogeny from Upham et al (2019) showing neonatal maturity data at tips and posterior probabilities of ancestral states at internal nodes. **(b)** Ancestral states at key nodes in the mammalian phylogeny. Posterior probabilities of neonatal maturity states at the given node are plotted on the Y-axis for each of the 100 sampled phylogenies represented on the X-axis. Xen = Xenarthra, Afroth = Afrotheria, Urs = Ursidae.

Surprisingly, the eutherian ancestor is reconstructed as precocial in all sampled trees but one (**Figure 1b**), with mean posterior probabilities 0.02 for altriciality and 0.98 for precociality. This result contradicts the prevailing assumption that the eutherian ancestor gave birth to altricial neonates.

## 4 Discussion

### 4.1 Sensitivity of ancestral state reconstruction

The ancestral state reconstruction described above is sensitive to taxon sampling. Data from Case (1978), biased toward boreoeutherian taxa, produce an altricial reconstruction of the eutherian ancestral state when used exclusively (**Supp. Figure 3**). However, expanding the taxonomic coverage shifts the estimated ancestral state toward precocial.

The reconstruction is also influenced by model choice. For Case (1978) data alone, the ER model is the best fit across all 100 trees, with mean model weights of 0.52 for ER, 0.33 for SYM, and 0.15 for ARD (**Supp. Figure 4**). Moreover, among the 100 trees, there is an apparent positive association between the ER model weight and the posterior probability of an altricial ancestral state for the eutherian node (**Supp. Figure 5a**). In the full dataset, the only tree for which the eutherian node is reconstructed as altricial also has the highest weight for the ER model among all trees (**Supp. Figure 5b**). Forcing character histories to be generated exclusively under the ER model with the full dataset reconstructs the eutherian ancestral state as altricial (**Supp. Figure 6**). Note, however, that the ER model provides the poorest fit for the full dataset.

In summary, using the full dataset with a balanced taxonomic sampling and following the model weights dictated by their fit strongly supports a precocial eutherian ancestral state.

### 4.2 Previous estimates of the eutherian ancestral state for neonatal maturity

The prevailing notion of an altricial eutherian ancestor stems, in part, from a long-held bias that marsupial reproductive strategies are a steppingstone toward more advanced eutherian strategies. Altriciality of the eutherian ancestor, as a result, is often assumed rather than inferred. This assumption is reinforced by the scattered distribution of precociality on eutherian phylogeny (Derrickson, 1992). However, this pattern may be illusory as the most speciose eutherian groups, such as Rodentia and Carnivora, tend to be altricial (**Supp. Table 1**).

The studies that have estimated the ancestral state using comparative methods tend to support an altricial eutherian ancestor, but their inference carries substantial uncertainty.

Werneburg et al (2016) estimated the eutherian ancestor’s age at eye opening and age at fur development to be 25 and 24 days after birth, respectively, suggesting an altricial condition. However, the authors note that the eutherian ancestor’s inferred values had the broadest confidence intervals, even exceeding those of older nodes. Notably, the 70% confidence intervals for eye opening and fur development overlap with those for gestation length (**Supp. Figure 7**). It cannot, thus, be conclusively inferred from these estimates that eye opening and fur development happened after birth in the eutherian ancestor.

O’Leary et al (2013) inferred that the eutherian ancestor had hairless neonates with closed eyes and a litter size of one. While hairlessness and closed eyes are associated with altriciality, the litter size of one is often associated with precocial neonates. The reconstruction suggests that these neonates, even if altricial, were not as altricial as the clearly altricial eutherians.

Ferner et al (2017) consider the possibility that both the eutherian and mammalian ancestors were precocial. However, they favor an altricial eutherian ancestor due to marginally better parsimony support, and similarly, an altricial mammalian ancestor based on prior evidence, such as the small size of Mesozoic mammals. While small mammals do tend to produce altricial neonates, it is not a necessary association. For instance, elephant shrew and spiny mouse are precocial, whereas bears and anteaters are altricial. The analysis by Ferner et al (2017) underscores the ambiguities inherent to inferring ancestral neonatal traits in mammals.

### 4.3 Evidence inconsistent with ancestral precociality

Our inference that the eutherian ancestor was precocial, likely requiring a long gestation, appears to conflict with the understanding of early eutherian reproductive physiology.

Eutherian pregnancy depends on sustained circulating progesterone levels, with the corpus luteum serving as the source early in gestation. However, eutherian gestation far exceeds the length of the luteal phase in a non-pregnant ovarian cycle. This extended, or “trans-cyclic” (Chavan et al., 2016), gestation is supported by a variety of “maternal recognition of pregnancy” mechanisms, which either extend the lifespan of the corpus luteum or activate an extra-ovarian source of progesterone, or both (Basanta & Pavlicev, 2025; Swaggart et al., 2015).

Divergent mechanisms of maternal recognition of pregnancy suggest that trans-cyclic gestation evolved independently in major eutherian lineages. That is, the eutherian ancestor had an intra-cyclic pregnancy (Basanta & Pavlicev, 2025; Chavan et al., 2016). A challenge with an ancestral intra-cyclic pregnancy is that extending its duration would also prolong the non-pregnant ovarian cycle. This is due to the luteal phase and pregnancy being serial homologs, which are expected to evolve in a correlated manner. Prolonged non-pregnant cycles would reduce the reproductive rate, posing a selective disadvantage (Basanta & Pavlicev, 2025). Through variational decoupling of the luteal phase from gestation, eutherian lineages could explore gestation lengths unbounded by the ovarian cycle. However, this decoupling was absent in the eutherian ancestor.

This leaves us with two seemingly incompatible results: (i) the eutherian ancestor gave birth to precocial neonates, likely requiring a long pregnancy, and (ii) ancestral eutherian pregnancy, being intra-cyclic, was constrained to a short duration that could not be extended without incurring a selective cost.

The resolution lies in the notion that the selective disadvantage of a long sterile ovarian cycle is not realized in species that have induced ovulation or breed seasonally (Basanta & Pavlicev, 2025). The eutherian ancestor is indeed inferred to have been an induced ovulator, with spontaneous ovulation evolving later (Pavlicev & Wagner, 2016). This ancestral condition would have allowed the eutherian stem lineages to prolong pregnancy without incurring the selective cost. Ancestral eutherian pregnancy could have been intra-cyclic yet long enough for the development of a precocial neonate.

### 4.4 Evidence consistent with ancestral precociality

Even if one does not accept our inference at face value (Griffith et al., 2015), ancestral precociality emerges as a plausible biological model in light of recent body of evidence.

#### 4.4.1 Reproductive traits enabling prolonged pregnancy predate the eutherian ancestor

Giving birth to precocial neonates requires a long pregnancy, a hallmark of eutherian mammals made possible by several evolutionary innovations. The evolution of embryo implantation (Griffith et al., 2017), facilitated by the modification of maternal inflammatory reaction (Chavan et al., 2017; Stadtmauer & Wagner, 2019) through the origin of decidual stromal cells (Chavan et al., 2020; Erkenbrack et al., 2018), enabled the establishment of a sustained fetal-maternal interface. The timing of these evolutionary events has been traced to the stem eutherian lineage, suggesting that the eutherian ancestor’s reproductive biology could support prolonged pregnancies.

#### 4.4.2 Precociality evolved early in mammalian evolution

Funston et al (2022) reconstructed the life history of an early Paleocene mammal *Pantolambda bathmodon* using paleohistology and geochemical analysis of dentine and enamel. Its reproductive mode resembled that of extant precocial eutherians, with the estimated gestation length of 207 days and a short suckling period of 31–56 days. Although the precise phylogenetic placement of *Pantolambda* remains unresolved (either related to Cetartiodactyla or a stem-group eutherian), it is clearly an early eutherian lineage (~62 million years old), providing evidence for precocial life history traits arising early in eutherian evolution.

Histology of long-bone fossils of multituberculates — a stem therian lineage — suggests that they had prolonged prenatal growth and abbreviated postnatal growth (Weaver et al., 2022), a pattern similar to eutherians but not marsupials. The patterns of cranial development across extant mammals and their ancestral state reconstruction also show that the ancestral therian condition resembled eutherian mammals more than marsupials (White et al., 2023, 2024). These studies suggest that the extreme altriciality of marsupials is a derived trait (Renfree, 1993; Smith, 2001; Smith & Keyte, 2020), and raise the possibility that eutherian-like reproductive traits — long intrauterine development and precocial neonates — may have evolved early in therian evolution.

### 4.5 A model for the evolutionary history of eutherian precociality

Our results and other evidence lead us to propose an alternative model for the evolutionary history of eutherian precociality, departing from the traditional narrative in three ways: (i) precociality arose early in therian evolution, (ii) marsupial altriciality is a derived trait, and (iii) eutherian ancestor likely gave birth to precocial neonates.

Precociality could have evolved early in mammalian phylogeny in two contexts: either in an oviparous mode prior to the mammalian ancestor, or in a viviparous mode in the therian stem lineage. If the latter, viviparity was likely achieved via egg retention. In this mode, a strategy observed in many squamate lineages (Blackburn, 2006), the egg is retained in the maternal reproductive tract until the embryo is near hatching, allowing extended gestation and precocial development. Crucially, the presence of an eggshell separates fetal and maternal tissues, preventing the maternal inflammatory reaction that would trigger early parturition (Wagner et al., 2014).

The viviparity seen in crown-group Theria likely evolved later along the therian stem lineage via early hatching in utero (Wagner et al., 2014). In this mode, the early loss of the eggshell or the zona pellucida leads to physical contact between fetal membranes and uterine endometrium causing an attachment-induced inflammatory reaction, precipitating parturition. This mechanism, culminating in birth shortly after early hatching, inherently resulted in extreme altriciality. We propose that this highly altricial state, derived from more precocial early therian lineages, became the ancestral condition for crown-group Theria and was subsequently retained and exaggerated in the marsupial lineage. Marsupials, in turn, evolved a suite of compensatory mechanisms to support adequate postnatal development (Smith & Keyte, 2020).

In the eutherian stem lineage, the inflammatory response from embryo attachment was modified into implantation (Griffith et al., 2017) through the evolution of decidual cells (Chavan et al., 2020). This allowed for the evolution of prolonged gestation and a re-emergence of precociality in the eutherian stem lineage. An indirect support for precociality in the eutherian stem lineage is provided by the observation that even the most altricial eutherian mammals are more developed at birth than the least altricial marsupials (Ferner et al., 2017; Halley, 2017; Szdzuy & Zeller, 2009; **Figure 2**). This implies that eutherian altriciality is not a direct continuation of the extreme altriciality of the therian ancestor but derived from a precocial intermediate state in the eutherian stem. Given its substantial length (60–90 million years), there were plausibly multiple transitions between precociality and altriciality along the eutherian stem lineage. The developmental state of the ancestral eutherian neonates — whether precocial or secondarily altricial (thus still more developed than marsupial neonates) — then depends on the most recent transition immediately preceding the crown-group Eutheria node (**Figure 3**).

**Figure 2:**
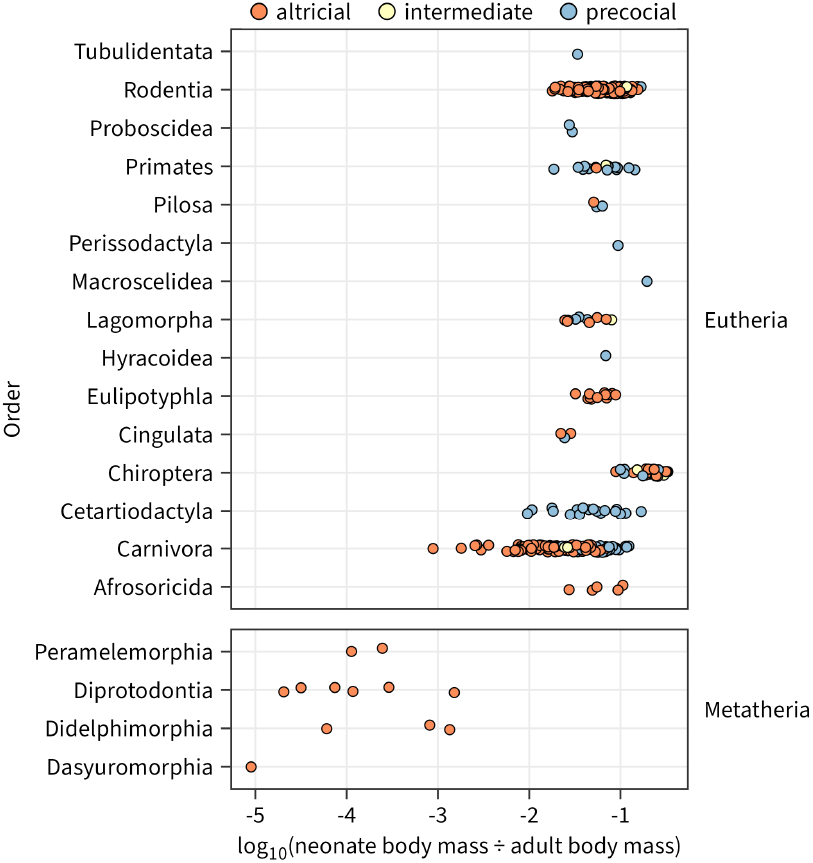
Altricial eutherian neonates are more developed than marsupials. Species with available adult and neonatal body mass data in PanTHERIA (Jones et al., 2009) are used. Log_10_-transformed relative neonatal body mass (RNBM) is used as a proxy for developmental maturity at birth. Higher the RNBM, greater the degree of maturity. Median log_10_(RNBM) for altricial eutherians and marsupials is –1.30 and –3.94, respectively, a difference of more than two orders of magnitude.

**Figure 3:**
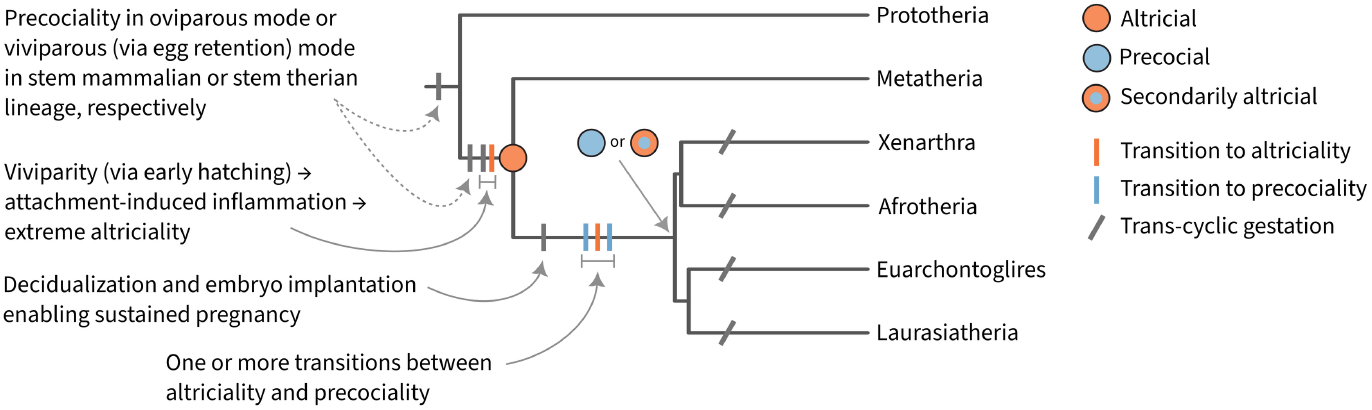
Model for the evolution of precociality in mammals. The key elements are an early evolution of precociality in mammalian phylogeny, reversal to altriciality in the therian stem lineage driven by early-hatching and attachment-induced inflammation, and re-evolution of precociality in the eutherian stem lineage enabled by the origin of decidual cells and embryo implantation.

Additional fossil data closer to the eutherian ancestor node will be necessary for a more direct evaluation of models of neonatal maturity. In the meantime, we encourage consideration of a precocial ancestry as a plausible scenario in eutherian evolution.

Our proposed model deviates from the simplistic narrative of mammalian reproductive evolution as a linear progression from an assumed primitive state to the derived states observed today. We believe that mammalian reproductive evolution is inherently non-parsimonious, shaped by an interplay of life history trade-offs, genomic conflict, pleiotropy among reproductive traits, anatomical and physiological constraints, and the origin of novel cell types and organs that break ancestral constraints and reshape the variational landscape. A deeper mechanistic understanding of these underlying processes and their evolution is critical for making sense of the complex evolutionary history of mammalian reproduction.

## 5 Methods

### 5.1 Data

Neonatal maturity data were collected from two sources: Case (1978) and PanTHERIA (Jones et al., 2009). Case (1978, Appendix 2) provides data on the developmental maturity at birth for 150 mammals scored on seven traits: body hair present, teeth present, open eyes, open auditory meatus, the ability to right itself when lying on its back, the ability to move forward, and formed claws. Species meeting at least 6 of the 7 criteria were classified as “precocial”, while those that did not meet at least 6 criteria were classified as “altricial”. Intermediate states were classified as “semi-altricial” or “semi-precocial”. Extremely altricial marsupials and monotremes were classified as “fetal”. We simplified this classification to three categories: altricial (fetal, altricial, semi-altricial), intermediate (semi-precocial), and precocial (precocial).

The variables “age at eye opening” and “litter size” from PanTHERIA were used to classify species by neonatal maturity status. All monotremes in the dataset had missing values for age at eye opening. After filling in the missing values for monotremes, 348 species (338 eutherians, 8 marsupials, 2 monotremes) had data for both variables. We followed the criteria of Martin and McLarnon (1985) with some modifications. Species with an age at eye opening more than 5 days were classified as altricial regardless of their litter size since this is a direct measure of maturity. Species with eyes open at birth and litter size under 1.5 were classified as precocial. Species with age at eye opening between 0 and 5 days, and litter size between 1.5 and 3 were classified as intermediates. All other species falling outside these bounds were manually classified based on literature.

The union of these two datasets was used as the final dataset. For species that are represented in both datasets, classification from Case (1978) was preferred over PanTHERIA. This dataset was filtered to remove species not represented in the phylogeny used for the analysis. The final dataset contains 420 species: 293 altricial, 115 precocial, and 12 intermediate (**Supp. Table 1**). The taxonomic coverage of all mammalian orders in the dataset can be found in **Supp. Table 2**.

### 5.2 Phylogeny

Phylogenetic framework from Upham et al (2019) was used for ancestral state reconstruction. We downloaded a sample of 100 trees from a credible set of 10,000 trees that capture uncertainties in inferences of tree topology and branch lengths (“Mammals birth-death node-dated DNA-only trees (4098 species, set of 10k trees)”), from http://vertlife.org/phylosubsets/. The trees were pruned to the subset of species present in the neonatal maturity dataset. The consensus phylogeny was also downloaded for visualization.

### 5.3 Ancestral state reconstruction

Ancestral state reconstruction was performed using stochastic character mapping (Bollback, 2006; Huelsenbeck et al., 2003; Revell, 2025) with ‘phytools’ version 2.1.1 (Revell, 2024) in R version 4.2.3 (R Core Team, 2023). The reconstruction was run on all 100 trees sampled from the credible set to account for phylogenetic uncertainty as well as on the consensus tree. Results from the latter were used for visualization of ancestral states on the phylogeny.

We fit three models using the ‘phytools::fitMk()’ function: equal rates (ER), symmetrical (SYM), and all rates different (ARD). We compared the model fits using ‘stats::anova()’. To account for model uncertainty, we generated 1,000 stochastic character histories using all three models in proportion of their model weights with the function ‘phytools::simmap()’. Posterior probabilities of ancestral states at all nodes were calculated by summing over all 1,000 stochastic character histories with the function ‘base::summary()’.

## Supporting information

Neonatal Maturity Data

Phylogenies

## 6 Data and code availability

Neonatal maturity data and the sample of 100 phylogenies from Upham et al (2019) used in this study are available as supplementary files. Code to reproduce all analyses is available at https://github.com/archavan/precocity.

## 7 Acknowledgements

We thank D. Stadtmauer, M. Srivastav, and I. Dhingra for their critical reading of an earlier version of this manuscript.

## References

Basanta, S., & Pavlicev, M. (2025). The Shifting Role and Regulation of the Corpus Luteum in Vertebrate Reproduction: A Synthetic Review. The Quarterly Review of Biology, 100(3), 189–231. 10.1086/737357

Blackburn, D. G. (2006). Squamate Reptiles as Model Organisms for the Evolution of Viviparity. Herpetological Monographs, 20, 131–146.

Bollback, J. P. (2006). SIMMAP: Stochastic character mapping of discrete traits on phylogenies. BMC Bioinformatics, 7(1), 88. 10.1186/1471-2105-7-88

Case, T. J. (1978). On the evolution and adaptive significance of postnatal growth rates in the terrestrial vertebrates. The Quarterly Review of Biology, 53(3), 243–282. 10.1086/410622

Chavan, A. R., Bhullar, B.-A. S., & Wagner, G. P. (2016). What was the ancestral function of decidual stromal cells? A model for the evolution of eutherian pregnancy. Placenta, 40, 40–51. 10.1016/j.placenta.2016.02.012

Chavan, A. R., Griffith, O. W., Stadtmauer, D. J., Maziarz, J., Pavlicev, M., Fishman, R., Koren, L., Romero, R., & Wagner, G. P. (2020). Evolution of embryo implantation was enabled by the origin of decidual stromal cells in eutherian mammals. Molecular Biology and Evolution, msaa274. 10.1093/molbev/msaa274

Chavan, A. R., Griffith, O. W., & Wagner, G. P. (2017). The inflammation paradox in the evolution of mammalian pregnancy: Turning a foe into a friend. Current Opinion in Genetics & Development, 47, 24–32. 10.1016/j.gde.2017.08.004

Danis, T., Lin, D., Caetano, D. S., Funston, G. F., & Rokas, A. (2025). Gestation length both shapes and is shaped by other life history traits in terrestrial eutherian mammals. Evolution Letters, qraf028. 10.1093/evlett/qraf028

Derrickson, E. M. (1992). Comparative reproductive strategies of altricial and precocial eutherian mammals. Functional Ecology, 6(1), 57. 10.2307/2389771

Erkenbrack, E. M., Maziarz, J. D., Griffith, O. W., Liang, C., Chavan, A. R., Nnamani, M. C., & Wagner, G. P. (2018). The mammalian decidual cell evolved from a cellular stress response. PLoS Biology, 16(8), e2005594. 10.1371/journal.pbio.2005594

Ferner, K., Schultz, J. A., & Zeller, U. (2017). Comparative anatomy of neonates of the three major mammalian groups (monotremes, marsupials, placentals) and implications for the ancestral mammalian neonate morphotype. Journal of Anatomy, 231(6), 798–822. 10.1111/joa.12689

Funston, G. F., dePolo, P. E., Sliwinski, J. T., Dumont, M., Shelley, S. L., Pichevin, L. E., Cayzer, N. J., Wible, J. R., Williamson, T. E., Rae, J. W. B., & Brusatte, S. L. (2022). The origin of placental mammal life histories. Nature, 610(7930), 107–111. 10.1038/s41586-022-05150-w

Griffith, O. W., Blackburn, D. G., Brandley, M. C., Van Dyke, J. U., Whittington, C. M., & Thompson, M. B. (2015). Ancestral state reconstructions require biological evidence to test evolutionary hypotheses: A case study examining the evolution of reproductive mode in squamate reptiles. Journal of Experimental Zoology Part B: Molecular and Developmental Evolution, 324(6), 493–503. 10.1002/jez.b.22614

Griffith, O. W., Chavan, A. R., Protopapas, S., Maziarz, J., Romero, R., & Wagner, G. P. (2017). Embryo implantation evolved from an ancestral inflammatory attachment reaction. Proceedings of the National Academy of Sciences, 201701129. 10.1073/pnas.1701129114

Grunstra, N. D. S., Zachos, F. E., Herdina, A. N., Fischer, B., Pavličev, M., & Mitteroecker, P. (2019). Humans as inverted bats: A comparative approach to the obstetric conundrum. American Journal of Human Biology, 31(2), e23227. 10.1002/ajhb.23227

Halley, A. C. (2017). Minimal variation in eutherian brain growth rates during fetal neurogenesis. Proceedings of the Royal Society B: Biological Sciences, 284(1854), 20170219. 10.1098/rspb.2017.0219

Huelsenbeck, J. P., Nielsen, R., & Bollback, J. P. (2003). Stochastic mapping of morphological characters. Systematic Biology, 52(2), 131–158. 10.1080/10635150390192780

Jones, K. E., Bielby, J., Cardillo, M., Fritz, S. A., O’Dell, J., Orme, C. D. L., Safi, K., Sechrest, W., Boakes, E. H., Carbone, C., Connolly, C., Cutts, M. J., Foster, J. K., Grenyer, R., Habib, M., Plaster, C. A., Price, S. A., Rigby, E. A., Rist, J., … Purvis, A. (2009). PanTHERIA: A species-level database of life history, ecology, and geography of extant and recently extinct mammals. Ecology, 90(9), 2648–2648. 10.1890/08-1494.1

Künkele, J., & Trillmich, F. (1997). Are Precocial Young Cheaper? Lactation Energetics in the Guinea Pig. Physiological Zoology, 70(5), 589–596. 10.1086/515863

Lillegraven, J. A. (1975). Biological considerations of the marsupial-placental dichotomy. Evolution, 29(4), 707. 10.2307/2407079

Lillegraven, J. A., Thompson, S. D., McNab, B. K., & Patton, J. L. (1987). The origin of eutherian mammals. Biological Journal of the Linnean Society, 32(3), 281–336. 10.1111/j.1095-8312.1987.tb00434.x

Martin, R. D., & MacLarnon, A. M. (1985). Gestation period, neonatal size and maternal investment in placental mammals. Nature, 313(5999), 220–223. 10.1038/313220a0

O’Leary, M. A., Bloch, J. I., Flynn, J. J., Gaudin, T. J., Giallombardo, A., Giannini, N. P., Goldberg, S. L., Kraatz, B. P., Luo, Z.-X., Meng, J., Ni, X., Novacek, M. J., Perini, F. A., Randall, Z. S., Rougier, G. W., Sargis, E. J., Silcox, M. T., Simmons, N. B., Spaulding, M., … Cirranello, A. L. (2013). The Placental Mammal Ancestor and the Post–K-Pg Radiation of Placentals. Science, 339(6120), 662–667. 10.1126/science.1229237

Pavlicev, M., & Wagner, G. P. (2016). The Evolutionary Origin of Female Orgasm. Journal of Experimental Zoology Part B: Molecular and Developmental Evolution, 326(6), 326– 337. 10.1002/jez.b.22690

Portmann, A. (1945). Die Ontogenese des Menschen als Problem der Evolutionsforschung. Verhandlungen Der Schweizerischen Naturforschenden Gesellschaft, 1, 44–53. 10.5169/seals-90447

R Core Team. (2023). R: a language and environment for statistical computing [Manual]. R Foundation for Statistical Computing. https://www.R-project.org/

Renfree, M. B. (1993). Ontogeny, Genetic Control, and Phylogeny of Female Reproduction in Monotreme and Therian Mammals. In F. S. Szalay, M. J. Novacek, & M. C. McKenna (Eds.), Mammal Phylogeny: Mesozoic Differentiation, Multituberculates, Monotremes, Early Therians, and Marsupials (pp. 4–20). Springer. 10.1007/978-1-4613-9249-1_2

Revell, L. J. (2024). phytools 2.0: An updated R ecosystem for phylogenetic comparative methods (and other things). PeerJ, 12, e16505. 10.7717/peerj.16505

Revell, L. J. (2025). Ancestral State Reconstruction of Phenotypic Characters. Evolutionary Biology. 10.1007/s11692-025-09645-y

Smith, K. K. (2001). The evolution of mammalian development. Bulletin of the Museum of Comparative Zoology, 156, 119–135.

Smith, K. K., & Keyte, A. L. (2020). Adaptations of the Marsupial Newborn: Birth as an Extreme Environment. The Anatomical Record, 303(2), 235–249. 10.1002/ar.24049

Stadtmauer, D. J., & Wagner, G. P. (2019). Cooperative Inflammation: The Recruitment of Inflammatory Signaling in Marsupial and Eutherian Pregnancy. Journal of Reproductive Immunology, 102626. 10.1016/j.jri.2019.102626

Starck, J. M., & Ricklefs, R. E. (Eds.). (1998). Avian growth and development: Evolution within the altricial-precocial spectrum. Oxford University Press.

Swaggart, K. A., Pavlicev, M., & Muglia, L. J. (2015). Genomics of Preterm Birth. Cold Spring Harbor Perspectives in Medicine, 5(2), a023127–a023127. 10.1101/cshperspect.a023127

Szdzuy, K., & Zeller, U. (2009). Lung and metabolic development in mammals: Contribution to the reconstruction of the marsupial and eutherian morphotype. Journal of Experimental Zoology Part B: Molecular and Developmental Evolution, 312B(6), 555–578. 10.1002/jez.b.21228

Upham, N. S., Esselstyn, J. A., & Jetz, W. (2019). Inferring the mammal tree: Species-level sets of phylogenies for questions in ecology, evolution, and conservation. PLOS Biology, 17(12), e3000494. 10.1371/journal.pbio.3000494

Wagner, G. P., Kin, K., Muglia, L., & Pavlicev, M. (2014). Evolution of mammalian pregnancy and the origin of the decidual stromal cell. The International Journal of Developmental Biology, 58(2–4), 117–126. 10.1387/ijdb.130335gw

Weaver, L. N., Fulghum, H. Z., Grossnickle, D. M., Brightly, W. H., Kulik, Z. T., Mantilla, G. P. W., & Whitney, M. R. (2022). Multituberculate Mammals Show Evidence of a Life History Strategy Similar to That of Placentals, Not Marsupials. The American Naturalist, 200(3), 383–400. 10.1086/720410

Werneburg, I., Laurin, M., Koyabu, D., & Sánchez-Villagra, M. R. (2016). Evolution of organogenesis and the origin of altriciality in mammals. Evolution & Development, 18(4), 229–244. 10.1111/ede.12194

White, H. E., Tucker, A. S., Fernandez, V., Miguez, R. P., Hautier, L., Herrel, A., Urban, D. J., Sears, K. E., & Goswami, A. (2023). Pedomorphosis in the ancestry of marsupial mammals. Current Biology, 33(11), 2136-2150.e4. 10.1016/j.cub.2023.04.009

White, H. E., Tucker, A. S., & Goswami, A. (2024). Divergent patterns of cranial suture fusion in marsupial and placental mammals. Zoological Journal of the Linnean Society, 203(2), zlae060. 10.1093/zoolinnean/zlae060

Zeveloff, S. I., & Boyce, M. S. (1980). Parental Investment and Mating Systems in Mammals. Evolution, 34(5), 973–982. 10.2307/2408002

