## Supplementary Text and Figures for "Precocial ancestry of placental mammals"

### Number of species and taxonomic coverage

Supp. Table 1: Number of species from mammalian orders used in the analysis and their neonatal-maturity.

| | | Altricial | Precocial | Intermediate |
|---|---|---|---|---|
| Prototheria | Monotremata | 2 | . | . |
| Metatheria | Dasyuromorphia | 1 | . | . |
| | Diprotodontia | 6 | . | . |
| | Peramelemorphia | 2 | . | . |
| | Didelphimorphia | 3 | . | . |
| Eutheria →Xenarthra | Cingulata | 2 | 1 | . |
| | Pilosa | 1 | 2 | . |
| Eutheria →Afrotheria | Afrosoricida | 5 | . | . |
| | Hyracoidea | . | 1 | . |
| | Macroscelidea | . | 1 | . |
| | Proboscidea | . | 2 | . |
| | Tubulidentata | . | 1 | . |
| Eutheria →Laurasiatheria | Carnivora | 86 | 34 | 3 |
| | Cetartiodactyla | . | 21 | . |
| | Chiroptera | 19 | 10 | 3 |
| | Eulipotyphla | 13 | . | . |
| | Perissodactyla | . | 2 | . |
| Eutheria →Euarchontoglires | Lagomorpha | 5 | 4 | 1 |
| | Primates | 1 | 13 | 1 |
| | Rodentia | 146 | 23 | 4 |
| | Scandentia | 1 | . | . |
| | Total | 293 | 115 | 12 |

Supp. Table 2: Taxonomic coverage of mammalian orders in the datasets used for analyses.

| | | Coverage (%) | | |
|---|---|---|---|---|
| | | Case 1978 | PanTHERIA | Combined |
| Prototheria | Monotremata | 20.00 | 40.00 | 40.00 |
| Metatheria | Dasyuromorphia | . | 1.41 | 1.41 |
| | Diprotodontia | 2.10 | 2.10 | 4.20 |
| | Microbiotheria | . | . | . |
| | Notoryctemorphia | . | . | . |
| | Peramelemorphia | . | 9.52 | 9.52 |
| | Didelphimorphia | 1.15 | 2.30 | 3.45 |
| | Paucituberculata | . | . | . |
| Eutheria ⟶Xenarthra | Cingulata | . | 14.29 | 14.29 |
| | Pilosa | 10.00 | 30.00 | 30.00 |
| Eutheria ⟶Afrotheria | Afrosoricida | 5.88 | 7.84 | 9.80 |
| | Macroscelidea | . | 6.67 | 6.67 |
| | Tubulidentata | . | 100.00 | 100.00 |
| | Hyracoidea | 25.00 | 25.00 | 25.00 |
| | Proboscidea | 33.33 | 66.67 | 66.67 |
| | Sirenia | . | . | . |
| Eutheria ⟶Laurasiatheria | Carnivora | 10.45 | 41.46 | 42.86 |
| | Cetartiodactyla | 6.15 | 0.31 | 6.46 |
| | Chiroptera | 0.81 | 2.69 | 2.87 |
| | Eulipotyphla | 0.67 | 2.92 | 2.92 |
| | Perissodactyla | 11.76 | 5.88 | 11.76 |
| | Pholidota | . | . | . |
| Eutheria ⟶Euarchontoglires | Lagomorpha | 6.52 | 7.61 | 10.87 |
| | Rodentia | 2.31 | 6.55 | 7.55 |
| | Dermoptera | . | . | . |
| | Primates | 3.72 | 0.53 | 3.99 |
| | Scandentia | 5.00 | . | 5.00 |

### Altriciality and precociality are distinct reproductive strategies

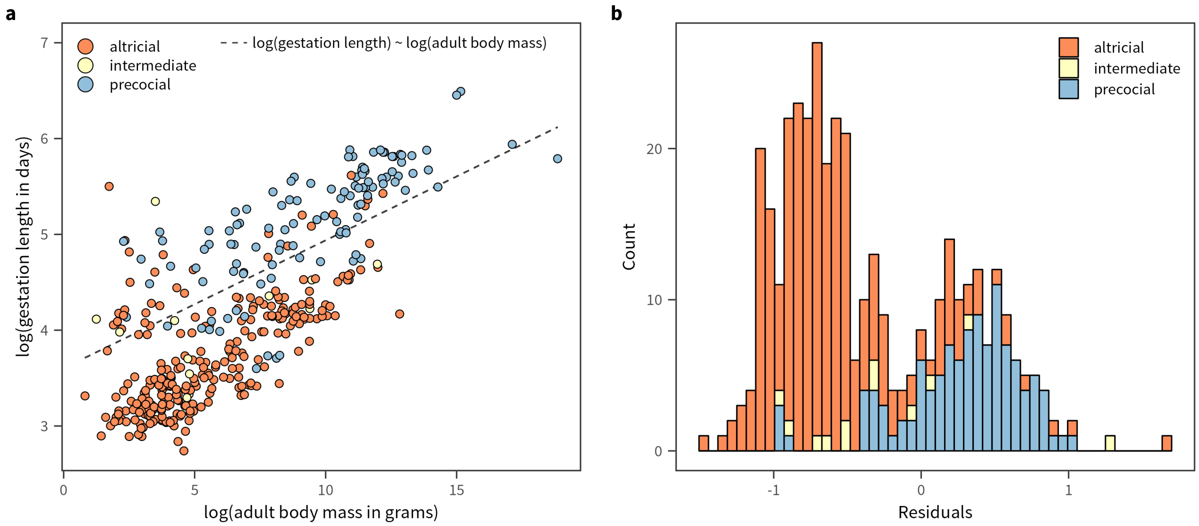


Supp. Figure 1: Altriciality and precociality are distinct life history strategies. This figure is a recreation of figures 1 and 2 from Martin & MacLarnon (1985) using the precocity data compiled in this study. **(a)** Phylogenetic generalized least squares (PGLS) (Grafen, 1989; Paradis & Schliep, 2018; Pinheiro et al., 2024) regression was performed using adult body mass as the predictor and gestation length as the response variable. The regression was run using the consensus phylogeny and was restricted to placental mammals for which adult body mass and gestation length data are available in PanTHERIA (Jones et al., 2009). **(b)** Histogram of the residuals of the model is plotted. After accounting for adult body mass, gestation length shows a bimodal distribution, with the two modes corresponding to altricial and precocial species.

### Ancestral state reconstruction

Supp. Table 3: Weights and AIC values for the three models used for ancestral state reconstruction using stochastic character mapping. The values here are for the consensus tree used for visualization in Figure 1 (main text). The model with best fit to the data is ARD.

| Model | Log Lik. | D.F. | AIC | Weight |
|---|---|---|---|---|
| ER | -153.3 | 1 | 308.6 | 0.002 |
| SYM | -150.144 | 3 | 306.289 | 0.005 |
| ARD | -141.943 | 6 | 295.886 | 0.993 |

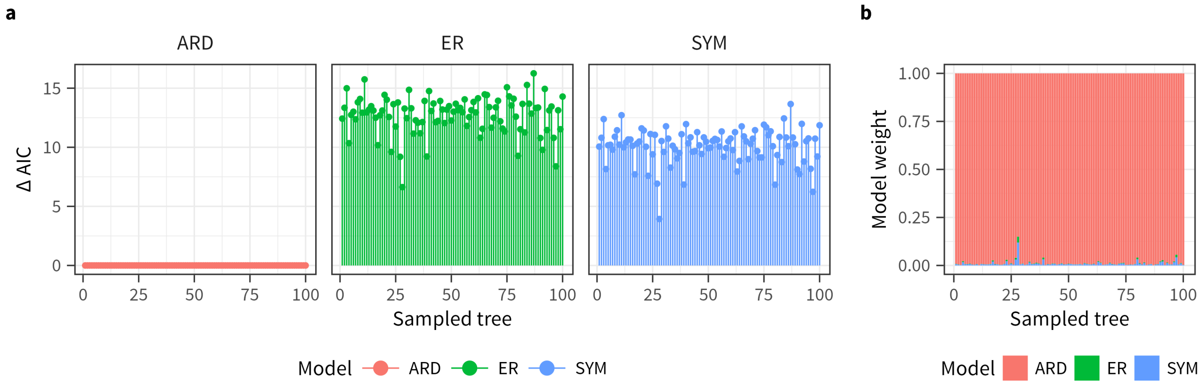


Supp. Figure 2: Δ AIC values **(a)** and weights **(b)** are shown for the three models used in ancestral state reconstruction via stochastic character mapping. The analysis was performed on a sample of 100 phylogenetic trees (Upham et al., 2019) (X-axis) to account for phylogenetic uncertainty. For each tree, 1,000 stochastic character histories were simulated based on the three models (Revell, 2024, 2025), with the proportion of histories assigned to each model determined by its weight. The best fit model, and thus with the highest weight, in all sampled trees is ARD.

### Effect of taxon sampling on the ancestral state reconstruction

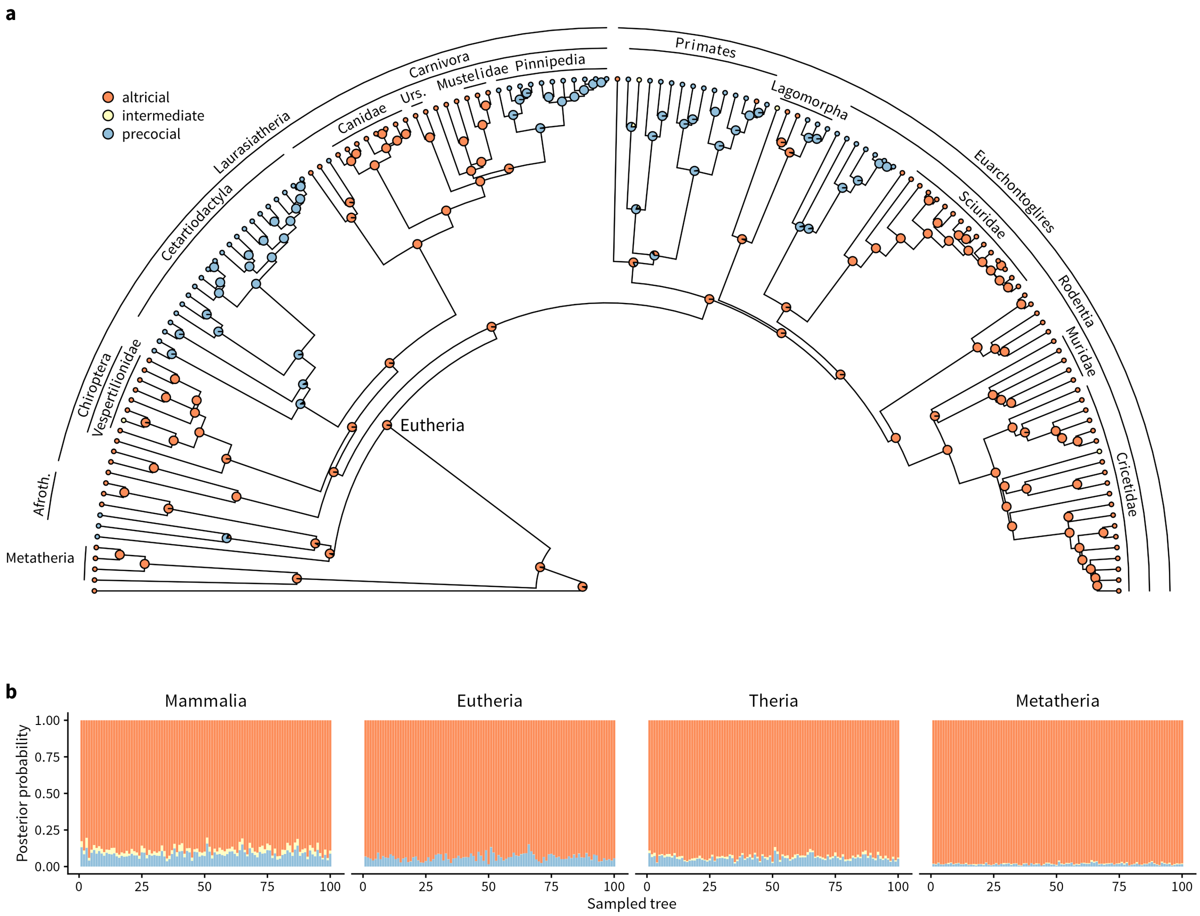


Supp. Figure 3: Effect of taxon sampling on ancestral state reconstruction. Ancestral state reconstruction of precocity was performed using species restricted to those from Case (1978). **(a)** Ancestral state reconstruction results plotted on the consensus phylogeny for visualization. **(b)** Posterior probabilities of ancestral states at four key nodes are shown, based on analyses run on a sample of 100 phylogenetic trees. For this subset of taxa, the reconstructed ancestral state for the crown-Eutherian node is consistently altricial across all 100 trees.

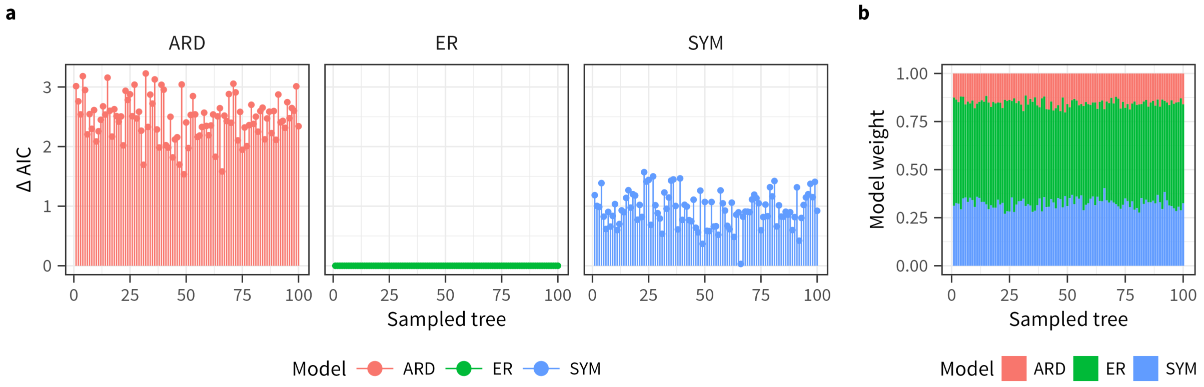


Supp. Figure 4: Δ AIC values **(a)** and weights **(b)** for models in the analyses shown in Supp. Figure 3 using Case (1978) subset of data. The model with lowest AIC consistently across all 100 trees was ER. However, note that the differences in AIC values for the three models in Supp. Figure 2 are much larger than for this subset of data. Consequently, the weights are relatively evenly distributed across models.

### Effect of model choice on the ancestral state reconstruction

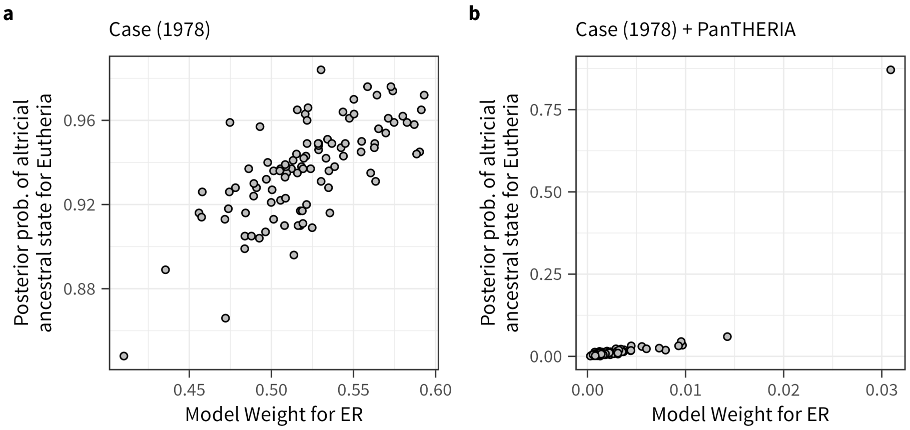


Supp. Figure 5: Effect of model choice on reconstructed ancestral state for precocity. Posterior probability for the reconstructed ancestral state being altricial for crown-Eutherian node across 100 sampled trees is plotted against the weight for ER model in those trees. **(a)** Results using the Case (1978) subset of the data. **(b)** Results using the full dataset, which includes both Case (1978) and PanTHERIA (Jones et al., 2009) data. In trees where the ER model has a higher weight, the posterior probability of an altricial ancestral state for the crown-Eutherian node tends to be higher.

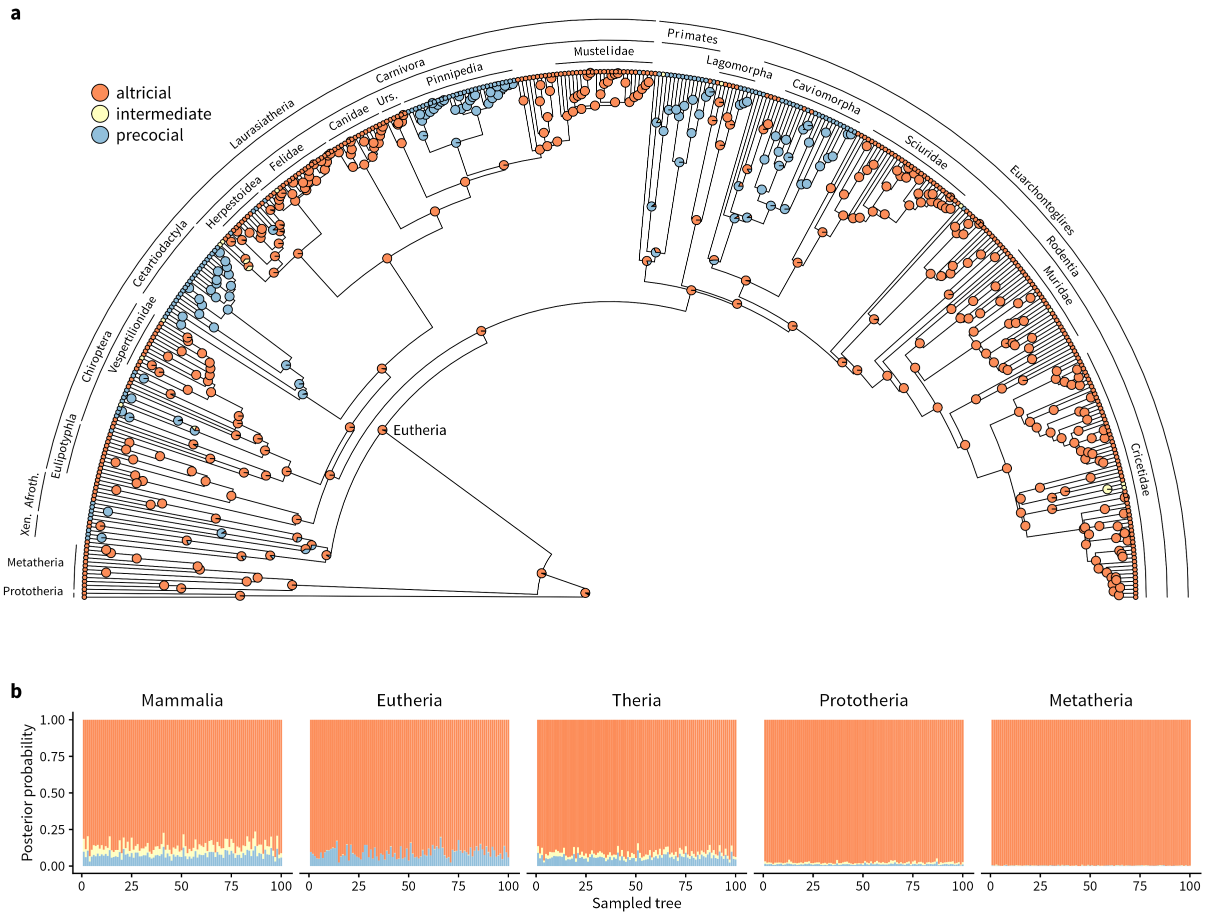


Supp. Figure 6: Effect of model choice on reconstructed ancestral state for precocity. Forcing the ER model for stochastic character mapping using the full dataset reconstructs the ancestral state for the crown-Eutherian node as altricial. Note, however, that the ER model is a poor fit for the full dataset; the best fitting model is ARD across all sampled 100 trees (Supp. Figure 2). **(a)** Reconstructed ancestral states plotted on consensus phylogeny for visualization. **(b)** Posterior probabilities for ancestral states at crown-Eutherian node and other key nodes across 100 sampled tree.

### Previous estimates of the eutherian ancestral state

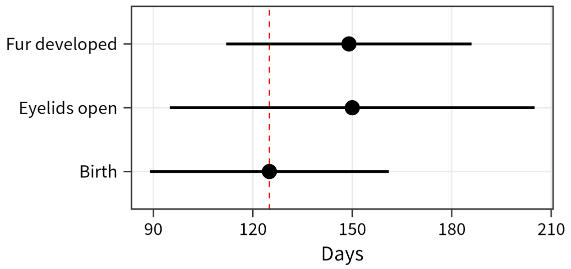


Supp. Figure 7: Ancestral state estimates of traits related to neonatal maturity for the crown-Eutherian node from Werneburg et al. (2016). Mean and 70% confidence intervals of days after conception are plotted on the X-axis (values from Table 1 in Werneburg et al). Even though the mean estimates for the age when the neonates open their eyes and their fur is developed are higher than the mean estimate for when they are born, the confidence intervals for the first two traits overlap with mean estimate for the latter (dotted red line).

### References


Case, T. J. (1978). On the evolution and adaptive significance of postnatal growth rates in the terrestrial vertebrates. *The Quarterly Review of Biology*, *53*(3), 243–282. https://doi.org/10.1086/410622

Grafen, A. (1989). The phylogenetic regression. *Philosophical Transactions of the Royal Society of London. B, Biological Sciences*, *326*(1233), 119–157. https://doi.org/10.1098/rstb.1989.0106

Jones, K. E., Bielby, J., Cardillo, M., Fritz, S. A., O'Dell, J., Orme, C. D. L., Safi, K., Sechrest, W., Boakes, E. H., Carbone, C., Connolly, C., Cutts, M. J., Foster, J. K., Grenyer, R., Habib, M., Plaster, C. A., Price, S. A., Rigby, E. A., Rist, J., … Purvis, A. (2009). PanTHERIA: a species-level database of life history, ecology, and geography of extant and recently extinct mammals. *Ecology*, *90*(9), 2648–2648. https://doi.org/10.1890/08-1494.1

Martin, R. D., & MacLarnon, A. M. (1985). Gestation period, neonatal size and maternal investment in placental mammals. *Nature*, *313*(5999), 220223. https://doi.org/10.1038/313220a0

Paradis, E., & Schliep, K. (2018). ape 5.0: an environment for modern phylogenetics and evolutionary analyses in R. *Bioinformatics*, *35*(3), 526–528. https://doi.org/10.1093/bioinformatics/bty633

Pinheiro, J., Bates, D., & R Core Team. (2024). *Nlme: Linear and nonlinear mixed effects models*. https://CRAN.R-project.org/package=nlme

Revell, L. J. (2024). phytools 2.0: an updated R ecosystem for phylogenetic comparative methods (and other things). *PeerJ*, *12*, e16505. https://doi.org/10.7717/peerj.16505

Revell, L. J. (2025). Ancestral State Reconstruction of Phenotypic Characters. *Evolutionary Biology*. https://doi.org/10.1007/s11692-025-09645-y

Upham, N. S., Esselstyn, J. A., & Jetz, W. (2019). Inferring the mammal tree: Species-level sets of phylogenies for questions in ecology, evolution, and conservation. *PLOS Biology*, *17*(12), e3000494.

https://doi.org/10.1371/journal.pbio.3000494

Werneburg, I., Laurin, M., Koyabu, D., & Sánchez-Villagra, M. R. (2016). Evolution of organogenesis and the origin of altriciality in mammals. *Evolution & Development*, *18*(4), 229244. https://doi.org/10.1111/ede.12194
